# Root vulnerability to embolism and lack of physiological recovery limit the competitive ability of an invasive palm

**DOI:** 10.1101/2025.07.30.667811

**Authors:** Thibaut Juillard, Christoph Bachofen, Maxwell Bergström, Timothy J. Brodribb, Hervé Cochard, Sylvain Delzon, Kate Johnson, Charlotte Grossiord

## Abstract

- Climate change and invasive species both threaten forest ecosystems. While exacerbated droughts drive tree mortality, species invasion alters forest composition. Invasive plants may be more vulnerable to drought due to acquisitive traits and low embolism resistance, but their potentially superior recovery capacity raises key uncertainties in predicting future species distributions.
- We compared the drought resistance and recovery of an invasive palm (*Trachycarpus fortunei*) with two native species (*Ilex aquifolium* & *Tilia cordata*) from the Southern Alps. Saplings were exposed to increasing intensities of drought-induced xylem embolism, then rewatered for 45 days. Among others, we tracked net assimilation (A_net_), stomatal conductance (g_s_), and leaf water potential (Ψ_leaf_) before and after the drought, and modeled the time to hydraulic failure in leaves, stems, and roots.
- While the invasive palm had exceptionally drought-tolerant leaves, its roots were the most drought-sensitive among all species. The palm maintained positive gas exchange until Ψ_leaf_ of -4 MPa, as some native species; however, it failed to recover from the most severe drought, with simulations showing root embolism preceding leaf failure, unlike native species.
- Invasive plants may match or exceed natives’ drought resistance, but their inability to recover from extreme droughts could limit their future spread.

## Introduction

Rising air temperatures and increasing drought frequency and severity are exacerbating forest mortality worldwide (Allen, Breshears and McDowell, 2015; Hammond *et al*., 2022). Simultaneously, forests face additional pressure from the spread of non-native plant species, which now affect nearly all ecosystems globally (Martin, Canham and Marks, 2009; Liebhold *et al*., 2017). These invasions alter species composition, reduce biodiversity, and disrupt ecosystem services (Peltzer *et al*., 2010; Liebhold *et al*., 2017).

As climate extremes intensify, understanding how invasive species respond to escalating drought stress has become increasingly urgent. Decreased precipitation could limit the spread of many invasive plants as they often have higher water requirements than native ones due to their resource-acquisitive traits (Cavaleri and Sack, 2010; Liu *et al*., 2017; Montesinos, 2022). Indeed, many invasive plants show consistently lower drought resistance than native ones at the leaf-level (Petruzzellis *et al*., 2021), yet can demonstrate similar or superior recovery after a drought event (Funk and Zachary, 2010; Barros *et al*., 2020; Zhang, Oduor and Liu, 2023). This paradox raises important but unresolved questions about how to assess drought vulnerability across species, especially when considering whole-plant responses and the role of individual organs. Crucially, drought recovery of perennial plants likely depends on the intensity of drought stress, particularly in relation to lethal thresholds, which remain largely misunderstood (Mcdowell, Brodribb and Nardini, 2019; Brodribb *et al*., 2021). Previous studies comparing the drought recovery of invasive and native plants using uniform drought durations found a better recovery in invasive plants (e.g., Puritty *et al*., 2019; Kelso, Wigginton and Grosholz, 2020; Zhang, Oduor and Liu, 2023), although it remains unclear whether these droughts were severe enough to cause meaningful hydraulic dysfunction. In contrast, Funk and Zachary (2010) applied drought based on relative water content to induce comparable stress levels across species and found no difference in recovery, underscoring the need for drought treatments calibrated to species-specific thresholds and functional assessments beyond leaves.

Drought resistance can be assessed from the vulnerability of the xylem to embolism (Choat *et al*., 2018), as the accumulation of emboli within the xylem leads to a rupture of the soil- atmosphere water continuum, ultimately resulting in the desiccation of plant organs, cell death, and plant mortality (McDowell *et al*., 2022; Mantova *et al*., 2023). Previous studies on embolism resistance (defined here as the water potential, *i*.*e*., Ψ, at which a given organ experienced a specific loss of xylem conductivity, typically 50% (P_50_)) in co-occurring invasive and native plants found mixed results: no difference (Pratt and Black, 2006), higher embolism resistance in invasive species (Crous, 2010; Yazaki *et al*., 2015), or the opposite (Zeballos *et al*., 2014; Petruzzellis *et al*., 2018), highlighting the need for a better understanding of hydraulic strategy differences between native and invasive plants. In addition, embolism resistance can differ between plant organs, with the most dispensable organs (*i*.*e*., leaves) typically having lower embolism resistance than more vital ones (*i*.*e*., roots and stems) (Johnson *et al*., 2016; Creek *et al*., 2018; Wilkening *et al*., 2023). This organ-level variation is rarely accounted for, despite offering critical insights into species’ drought survival strategies (Wilkening *et al*., 2023). For instance, species that show similar embolism resistance across organs typically produce costly and long-lived leaves and follow a resource-conservatism strategy (Wright *et al*., 2004; Smith-Martin *et al*., 2020). However, how these organ-level hydraulic traits vary between native and invasive species remains poorly understood, limiting our ability to predict the drought resilience and potential spread of invasive plants under increasing water stress.

Aside from embolism resistance, drought resistance is also mediated by stomatal regulation. While a decrease in stomatal conductance (g_s_) under drought decreases the risk of embolism, it also limits net assimilation (A_net_), potentially affecting the competitive ability of a species (Schönbeck *et al*., 2022). Invasive plants usually display higher g_s_ and carbon uptake than native ones, a pattern linked to greater water transport capacity and wider xylem vessels (Cavaleri and Sack, 2010; Díaz de León Guerrero *et al*., 2020; Petruzzellis *et al*., 2021). Therefore, while large vessel diameters could increase A_net_ under favorable conditions, they could also increase vulnerability to embolism during drought (Scoffoni *et al*., 2017). Yet it remains unclear whether invasive species modulate stomatal behavior differently under drought compared to natives, and how such regulation balances carbon gain with hydraulic risk.

Finally, damage to the photosynthetic apparatus itself - especially leaf dehydration - can impair recovery after rewatering (Brodribb and Cochard, 2009; Xu, Zhou and Shimizu, 2010; Trueba *et al*., 2019). This can be assessed via leaf chlorophyll fluorescence (F_v_/F_m_, *i*.*e*., the maximum quantum efficiency of Photosystem II), which closely tracks both tissue dehydration and the build-up of xylem embolism (Johnson, Jordan and Brodribb, 2018). Yet, no studies have combined embolism measurements, organ-level vulnerability, and photosynthetic recovery to characterize whole-plant drought resilience in invasive species.

Here, we address these gaps by comparing the drought resistance and recovery capacity of an invasive palm, *Trachycarpus fortunei*, with two co-occurring native species, *Ilex aquifolium* and *Tilia cordata*, growing in the southern Alps. While *T. fortunei* is currently expanding across many regions worldwide (Walther *et al*., 2007), its performance under increasingly dry climatic conditions remains uncertain, with potentially disastrous consequences for natural ecosystems if it proves both drought-tolerant and capable of rapid recovery from stress. As a palm, *T. fortunei* features a highly efficient water transport system enabled by broad xylem vessels (Olson *et al*., 2014). However, it lacks secondary growth and dormancy mechanisms (Renninger and N. Phillips, 2016), making it potentially more vulnerable to xylem embolism. On the other hand, the large amount of water stored in palm petioles and stems may buffer embolism formation and help sustain transpiration under mild drought stress (Emilio *et al*., 2019). We hypothesized that:

1. *T. fortunei* is less drought-resistant than the native species, as indicated by less negative P_50_, P_80_, and P_99_ at the leaf, petiole/stem, and root level, with leaves being the most vulnerable organs. Additionally, *T. fortunei* displays a more resource-acquisitive behavior characterized by high A_net_ and g_s_.
2. Once leaf P_50_, P_80,_ or P_99_ is reached, *T. fortunei* shows greater leaf tissue dehydration than native species (*i*.*e*., lower F_v_/F_m_) due to rapid embolism formation in the leaf xylem.
3. After rewatering, the recovery of *T. fortunei* depends on the drought severity: individuals exposed to leaf P_50_ recover better (higher A_net_ and F_v_/F_m_) than native species, thanks to more efficient water transport (higher g_s_). In contrast, plants exposed to more severe drought (P_80_ and P_99_) do not show improved recovery, as embolism is likely present in all organs, limiting the reactivation of hydraulic function.

## 2. Materials and Methods

### 2.1 Experimental set-up

In March 2023, we excavated 150 saplings of 3-4 years old (50-70 cm in height) *Trachycarpus fortunei* in a freshly invaded sub-Mediterranean forest (Cugnasco, Switzerland; 46°10’15’’ N, 8°55’49’’ E; 205 m a.s.l.; MAT: 12.2 °C; MAP: 1757 mm). At the same time, we bought 100 even-aged saplings of the native deciduous *Tilia cordata* and evergreen *Ilex aquifolium* from a commercial producer (Wiler bei Utzenstorf, CH). All plants were grown for one year under identical controlled conditions before the experiment, equalising trait acclimation to the same environmental conditions. All individuals were immediately potted in 7 L pots with generic sandy forest soil made of 20 % peat and 80 % mineral substrate (pH = 6.3; Ökohum; DE) and were stored at the Swiss Institute for the Forest, Landscape and Snow (WSL, 47°21’41.6“N, 8°27’21.5”E, Birmensdorf, 540 m a.s.l., MAT: 12.0 °C, MAP: 1008 mm). In October 2023, all plants were transported to the Swiss Federal Institute of Technology Lausanne (EPFL, 46°31’15.3“N, 6°34’04.0”E; Lausanne; 540 m a.s.l.; MAT: 12.6 °C; MAP: 1136 mm) and positioned randomly in two adjacent semi-automated plastic greenhouses with similar environmental conditions. Plants were watered at field capacity using a drip irrigation system equipped with individual pressure regulators (Allenspach GreenTech AG, CH) to ensure uniform irrigation for each pot. Air temperature (T_air_) and vapor-pressure deficit (VPD) were identical in the two greenhouses before and during the experiment (Fig. S1).

### 2.2 Embolism resistance of leaves, stems, and roots

To assess the embolism resistance of leaves, stems, and roots, we measured xylem embolism accumulation simultaneously in each plant organ of five to nine replicates of each three species in June 2024. Embolism in the three organs was tracked with the optical method as described in Brodribb *et al*. (2017) by drying out the entire sapling on a bench at a stable temperature and humidity. One ‘cavicam’ (cavicam.co, Hobart) was fixed on an undamaged leaf, stem, and root of each individual, and high-resolution images were automatically registered every 5 minutes. Embolism was quantified by computing the change in light transmission between two successive images using the optical vulnerability method (http://www.opensourceov.org) and the software ImageJ (NIH). Organs were considered fully embolized if no additional embolism event was observed within 6 hours. Simultaneously, the stem water potential (Ψ_stem_) was measured with psychrometers (PSY1 Psychrometer; ICT International, Australia). For *T*. *fortunei*, we measured the Ψ and xylem embolism of the petiole and not the pseudo-trunk (Fig. S2), which we refer to as “stem” in the following. For each organ, we fitted the cumulative embolism response to Ψ_stem_ with a logistic curve (>100 points per cumulative curve) (Fig. S3) and took Ψ_stem_ at 50, 80, and 99% of the total cumulated embolism to obtain P_50_, P_80_, and P_99_.

### 2.3 Drought-induced cell damage

In spare individuals of the three species, we quantified the drought-induced cell damage of every organ with measurements of relative electrolyte leakage (REL). For this, we removed 3-6 individuals per species from the pots and let them dry on a bench at a stable temperature and humidity while measuring Ψ_stem_ with psychrometers. Individuals were covered with dark plastic bags to prevent photosynthesis. Twice per day for two to three weeks, we collected leaf, stem, and root material from each individual, except in *T*. *fortunei*, where petioles were not sampled due to their low number (<10) per individual. Each cut on the plant was immediately sealed with melted candle wax to prevent localized plant dehydration and not resampled. Samples were then cleaned with deionized water and immediately placed in 50 ml Falcon tubes containing 20 ml of a solution of distilled water and 0.1% Triton X-100. After 24 hours of incubation at 4°C, the electrolyte leakage (EL) of each sample was measured with a conductivity probe (TetraCon 925; WTW, Germany) and a multimeter (Multi 3430, WTW, Germany). All samples were subsequently autoclaved at 120 °C for 20 minutes and measured again to obtain the maximum conductivity value (EL_max_). REL was then calculated by dividing EL by EL_max_. We fitted the REL response to Ψ_stem_ with a logistic curve (12-18 points per curve) and took the inflection point of the regression to obtain REL_50_ (*i*.*e*., the Ψ_stem_ at which REL of a given organ is at 50% of its maximum) (Fig. S4).

### 2.4 Photosynthetic and hydraulic traits during the drought and after rewatering

At the beginning of the experiment in June 2024 (T_0_), 18 fully hydrated individuals per species were exposed to extreme drought (no irrigation) until individuals reached the Ψ_leaf_ which induces leaf P_50_, P_80_, and P_99_ (T_1_; n = 6 for each intensity) that was determined by previous embolism resistance measurements (Fig. 1). We estimated the leaf water potential (Ψ_leaf_) from T_0_ to T_1_ on all individuals every 2-4 days using a Scholander-type pressure chamber (PMS Instruments; USA). To do so, we wrapped one leaf per individual in a plastic bag and aluminum foil around noon and waited for an hour to ensure stomatal closure was complete before measuring Ψ_leaf_. Some individuals of *T*. *cordata* lost most of their leaves before reaching P_50_, and we therefore measured their Ψ_stem_ using the same method to track the intensity of drought stress. Following this, plants were fully rewatered for 45 days to track their recovery (T_2_). Throughout the experiment, we monitored the soil water content using a TDR-100 (Spectrum Technologies, USA) to verify the effectiveness of the drought and rewatering manipulation (Fig. S5). Every two days, we measured leaf net assimilation (A_net_) and stomatal conductance (g_s_) with a photosynthetic system (Walz GFS-3000, Effeltrich, Germany) on three individuals per species and treatment. All measurements were taken under constant saturating light (PAR = 1500), T_air_ of 20°C, VPD of 1 kPa, and CO₂ concentration of 420 ppm. We let the gas exchange stabilize for at least 5 minutes before taking measurements. On an adjacent leaf, we measured the maximum quantum efficiency of Photosystem II (F_v_ /F_m_) with a FluorPen (Photon System Instruments; CZ). These leaves were wrapped in aluminum foil for 30 minutes before measurement to ensure dark acclimation. To assess drought resistance and resilience, we calculated the percentage of A_net_, g_s_, and F_v_/F_m_ at T_1_ and T_2_ relative to T_0_ for drought resistance and resilience, respectively.

**Fig. 1:**
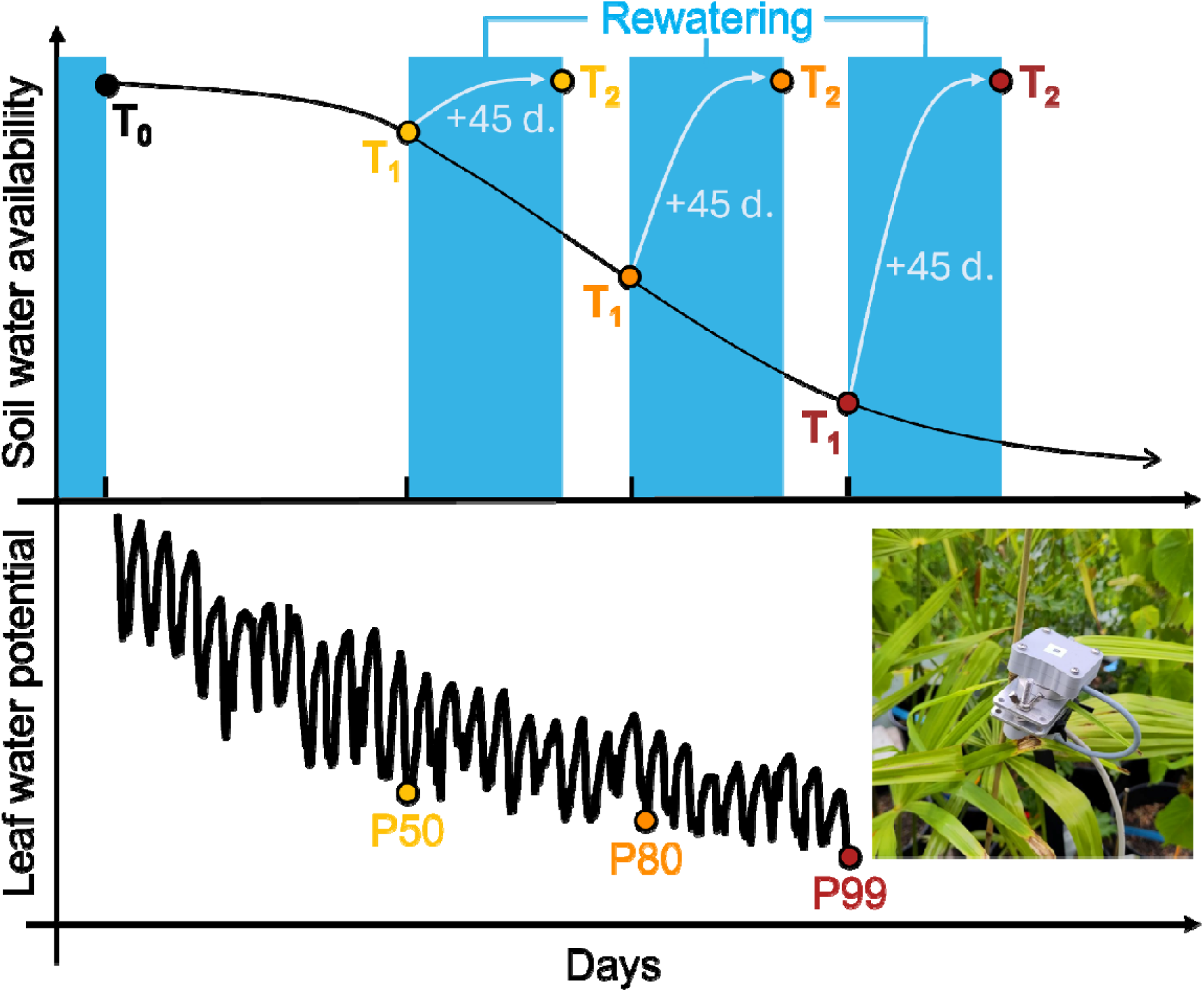
Schematic representation of the drought manipulation. Before the manipulation, potted plants of the three species (n = 18 plants per species) were watered to soil capacity (T_0_). The irrigation was then stopped until the leaf water potential of each individual reached P_50_, P_80_, and P_99_ (*i*.*e*., the leaf water potential at 50, 80, and 99% of leaf embolism, respectively) (T_1_) (n = 6 plants per species and drought intensity). Plants were then immediately rewatered to soil capacity and tracked for 45 days (T_2_). Leaf water potential was obtained from the leaf shrinkage measured optically (inserted picture), and scaled with water potential point measurements.

### 2.5 Time to hydraulic failure

To assess how the drought resistance of each organ influences the progression of hydraulic failure at the whole-plant level, and to evaluate species’ sensitivity to warming, we predicted the time to hydraulic failure (THF, defined here as P_99_) for each species using the SurEau model (Version 10/01/2025; Cochard *et al*., 2021). For this, we conducted 30 simulations per species under ambient (mean T_air_ = 20.7°C) and higher air temperature (*i*.*e*., +2.5, 5, 10, 15, and +20°C increase in mean T_air_). Trait responses to warming were extrapolated based on stomatal sensitivity to temperature, estimated from our measurements (Table S1). Each of the 30 simulations was parameterized with randomized inputs derived from measurements on five individuals per species. These included organ-specific vulnerability curves (P_50_ and slope at P_50_), gas exchange traits (*e*.*g*., maximum leaf conductance), and morphological attributes (*e*.*g*., stem diameter, height, and canopy height) (see Table S1 for all inputs). To improve the model accuracy, we estimated minimum leaf conductance (g_min_) using the detached leaf mass loss method (Pearcy, Schulze and Zimmermann, 2000) on four to five well-hydrated saplings per species. Before dawn, one branch (or leaflet for *T*. *fortunei*) per individual was sampled, and a single leaf per branch was sealed with wax at the petiole, scanned, weighed, and placed in a dark, climate-controlled room (19.9°C, 66% RH) monitored by a datalogger (HOBO MX2301A). Leaf mass was recorded every 15 minutes using a precision balance until at least 8 measurements captured the linear portion of the mass loss curve. g_min_ was then calculated as the transpiration rate divided by VPD. Similarly, we measured the leaf water potential at the turgor loss point (Ψ_TLP_) and elasticity modulus (ε) on four to five well-hydrated saplings per species using the pressure-volume curve method as described in Koide (2000). We also measured the leaf area index (LAI) of five individuals per species by calculating the total leaf area of scanned leaves using the software ImageJ (Schindelin *et al*., 2012) and dividing it by the projected crown area of each individual. The complete list of model inputs is provided in Table S1.

### 2.6 Statistical analyses

Species and organ differences in P_50_, P_80_, P_99_, and REL_50_ were determined through analysis of variance using species (*T*. *fortunei*, *I*. *aquifolium*, and *T*. *cordata*) and organs (leaf, stem, root) and their interaction as fixed effects. Similarly, statistical differences in drought resistance and resilience were calculated with species, drought treatments (P_50_, P_80_, and P_99_), and their interaction as fixed effects. Individual differences in P_50_, P_80_, P_99_, REL_50_, drought resistance, and drought recovery between species and organs (or species and drought treatments, respectively) were estimated with Tukey’s HSD post hoc tests. g_s_ and Ψ_leaf_ were log-transformed to ensure homoscedasticity. Finally, differences in THF between organs were assessed with Tukey’s HSD post hoc tests for each species and degree of warming separately. All analyses were executed with R software (4.3.2, R Core Team, 2021).

## 3. Results

### 3.1 Differences in embolism resistance and cell damage among species and organs

Overall, *T*. *fortunei* also showed a wider range of embolism resistance across tissues than both natives) (Fig. 2). Leaves had a much more negative P_50_ (-7.44 MPa) than roots (-1.33 MPa, *p* = 0.02) and, to a lesser extent, stems (-4.66 MPa, n.s.) (Fig. 2). Contrastingly, embolism resistance did not differ between organs in either native species (Fig. 2). The leaves of *T. fortunei* were also more drought-tolerant (-7.44 MPa) than the ones of *I*. *aquifolium* (-5.09 MPa, n.s.) and *T*. *cordata* (-2.41 MPa, *p* = 0.015).

**Fig. 2:**
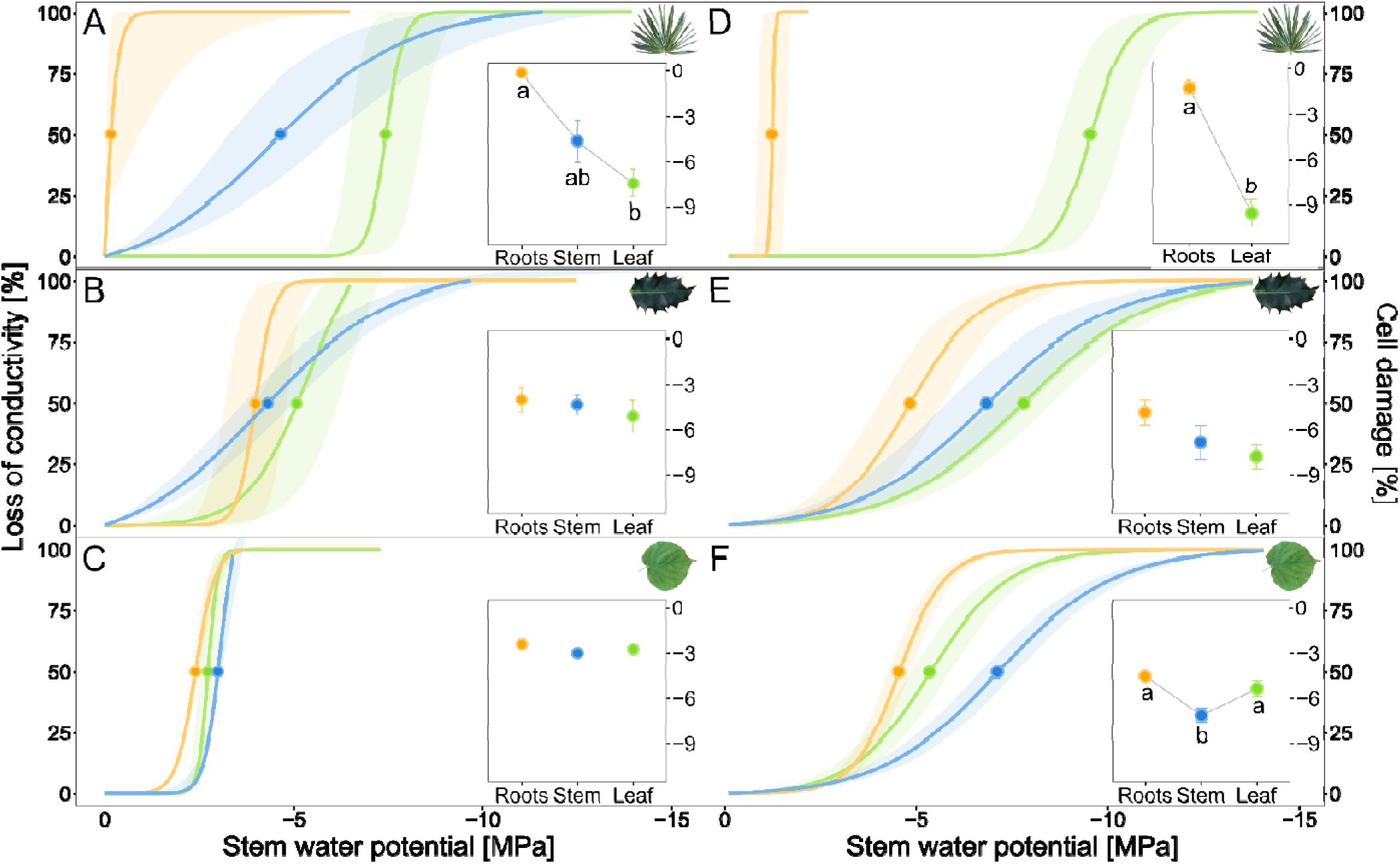
Loss of conductivity (A-C) and cell damages (D-F) in function of the stem water potential for each species (*T*. *fortunei*: A and D; *I*. *aquifolium*: B and E; *T*. *cordata*: C and F), and organs (green = leaves; blue = stems; orange = roots). For *T*. *fortunei*, petioles were measured instead of stems, but cell damage could not be assessed due to the presence of fewer than 10 petioles per individual. Means and standard errors are shown in thick and shaded colors, respectively (n = 3-9). Inserts show the mean and standard errors of conductivity loss or cell damage at 50%. Different letters indicate significant differences (p < 0.05) between organs.

P_50_ values were strongly correlated with REL_50_ (*p* = 0.001, R^2^ = 0.84; Fig. 3), which was consistently 2 MPa below P_50_. An exception was observed in the roots of *T*. *fortunei*, where the two metrics were nearly identical (-1.33 and -1.23 MPa for P_50_ and REL_50_, Fig. 3). REL_50_ further confirmed that roots of *T*. *fortunei* were less drought-tolerant than *I*. *aquifolium* (n.s) and *T*. *cordata* (*p* = 0.003), as indicated by their less negative REL_50_. Consistently across species and organs, stem P_50_ and REL_50_ values generally fell between those of leaves and roots, except stem in *T*. *cordata,* where the REL_50_ (-7.12 MPa) was more negative than the leaf (-5.36 MPa, *p* = 0.035) and root (-4.55 MPa, *p* = 0.003) (Fig. 2).

**Fig. 3:**
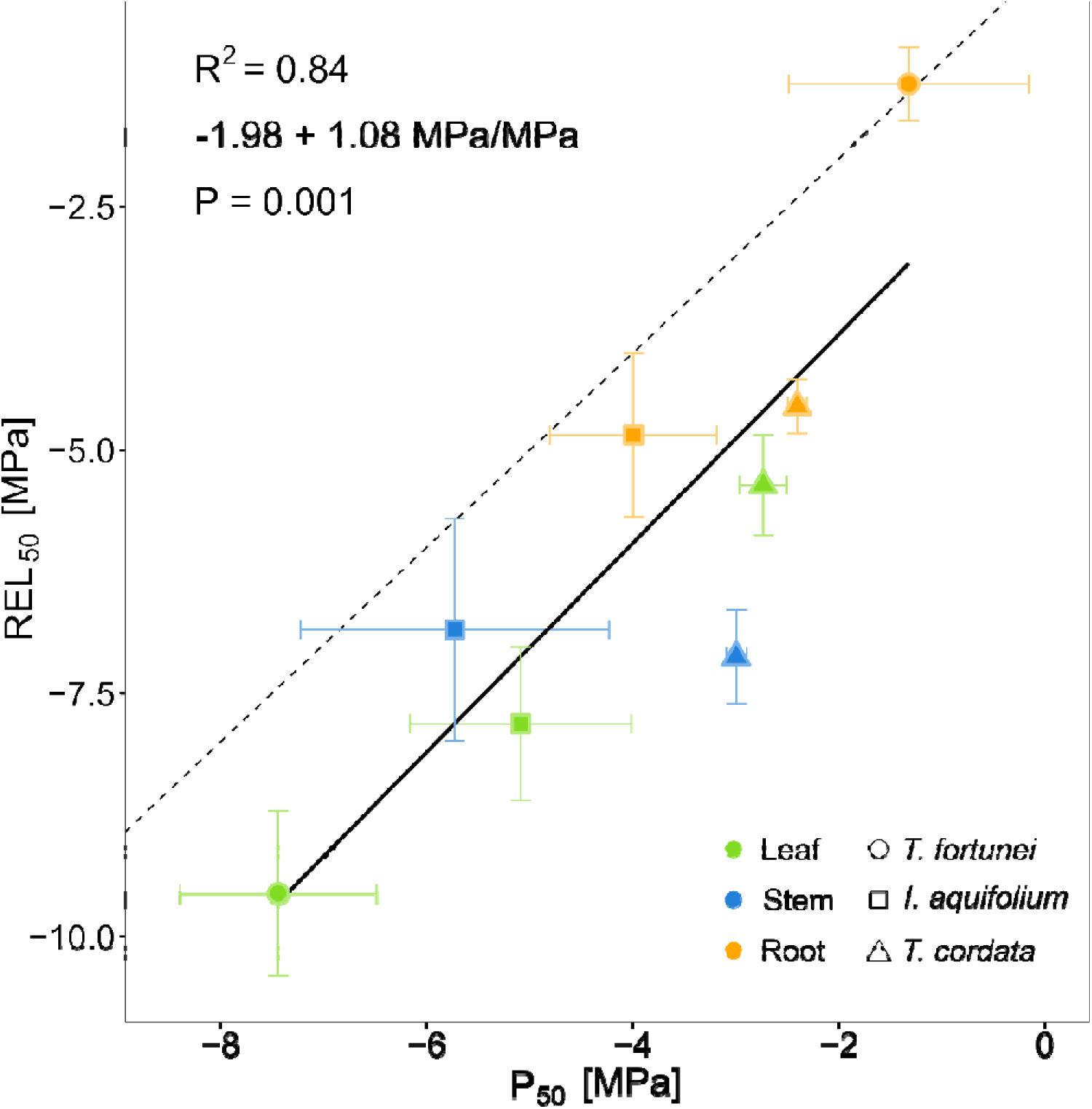
Stem water potential at 50% cell damage (REL_50_) in relation to the stem water potential at 50% loss of conductivity (P_50_) (mean ± s.e.) for each organ and species (n = 3-9). Colors represent the three organs, and shapes correspond to the three species. The 1:1 line is indicated with a dashed line.

### 3.2 Photosynthetic response to soil drought

At the beginning of the drought, when trees were well-watered, A_net_ and g_s_ were lower in *T*. *fortunei* than in *I*. *aquifolium* and *T*. *cordata* (A_net_ = 1.88 *vs*. 4.05 & 3.91 μmol m^2^ s^-1^, respectively, both *p* < 0.05; g_s_ = 14.6 *vs*. 71.6 & 41.1 mmol m^2^ s^-1^, *p* < 0.001 & *p* = 0.08, respectively). In all species, A_net_ and g_s_ decreased exponentially with the decreasing Ψ_leaf_ induced by the drought (R^2^ = 0.53, 0.29, and 0.60 for *T*. *fortunei*, *I*. *aquifolium*, and *T*. *cordata*, respectively; Fig. 4). All three species reached near-zero A_net_ and g_s_ around -2 MPa, yet *I*. *aquifolium* (and to a lower extent *T*. *fortunei*) showed occasionally positive values of A_net_ and g_s_ down to -4 MPa (Fig. 4. F_v_ /F_m_ was correlated with Ψ_leaf_ in all species (R^2^ = 0.64, 0.75, and 0.27 for *T*. *fortunei*, *I*. *aquifolium*, and *T*. *cordata*, respectively) (Fig. 4) and decreased linearly from 0.83 to 0 at a similar rate in all species (mean across species: -0.092 per MPa Ψ_leaf_). Importantly, in deciduous *T*. *cordata*, F_v_ /F_m_ dropped abruptly between 2 and 3 MPa due to leaf shedding (Fig. S5), while both invasive and native evergreen species kept some photosynthetically active leaves (F_v_ /F_m_ > 0) down to -7 MPa (Fig. 4).

**Fig. 4:**
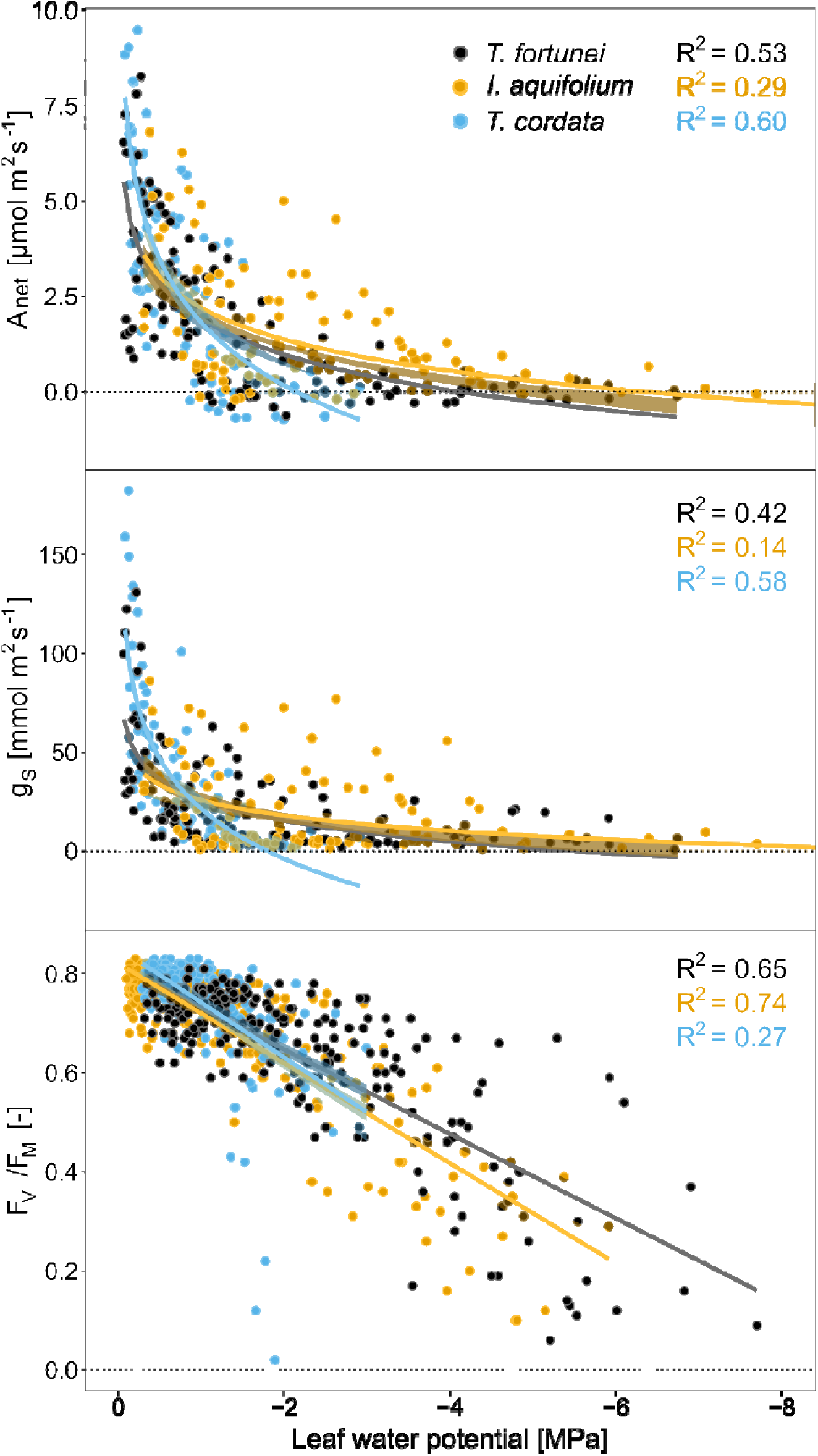
Net CO_2_ assimilation (A_net_), stomatal conductance (g_S_), and chlorophyll fluorescence (F_V_/F_M_) in relation to leaf water potential for each species. Regression lines were added when significant (P < 0.05).

The relative drop in A_net_ from T_0_ to T_1_ (A_net_ drought resistance) differed between species (*p* = 0.048) but not drought severity (n.s). *T*. *fortunei* was slightly less resistant than *I*. *aquifolium* (7.3% *vs*. 10.6% of the initial A_net_) but more so than *T*. *cordata*, which retained no measurable A_net_ (only one leaf left at 0% A_net_) (Fig. 5). Similarly, g_s_ resistance to drought varied between species (*p* = 0.01) and drought severity (*p* = 0.025), with *T*. *fortunei* maintaining markedly higher g_s_ at peak drought compared to the native species (67.4, 28.5, and 6.74 % of the initial g_s_, respectively), and being less impacted under P_50_ than more extreme drought levels (P_80_ and P_99_) (Fig. 5). F_v_ /F_m_ resistance varied between species *p* = 0.004), with *T*. *fortunei* experiencing a larger decline after the drought than *I*. *aquifolium* and *T*. *cordata* (13.0, 21.4, and 13.8%, respectively) (Fig. 5 & S6).

**Fig. 5:**
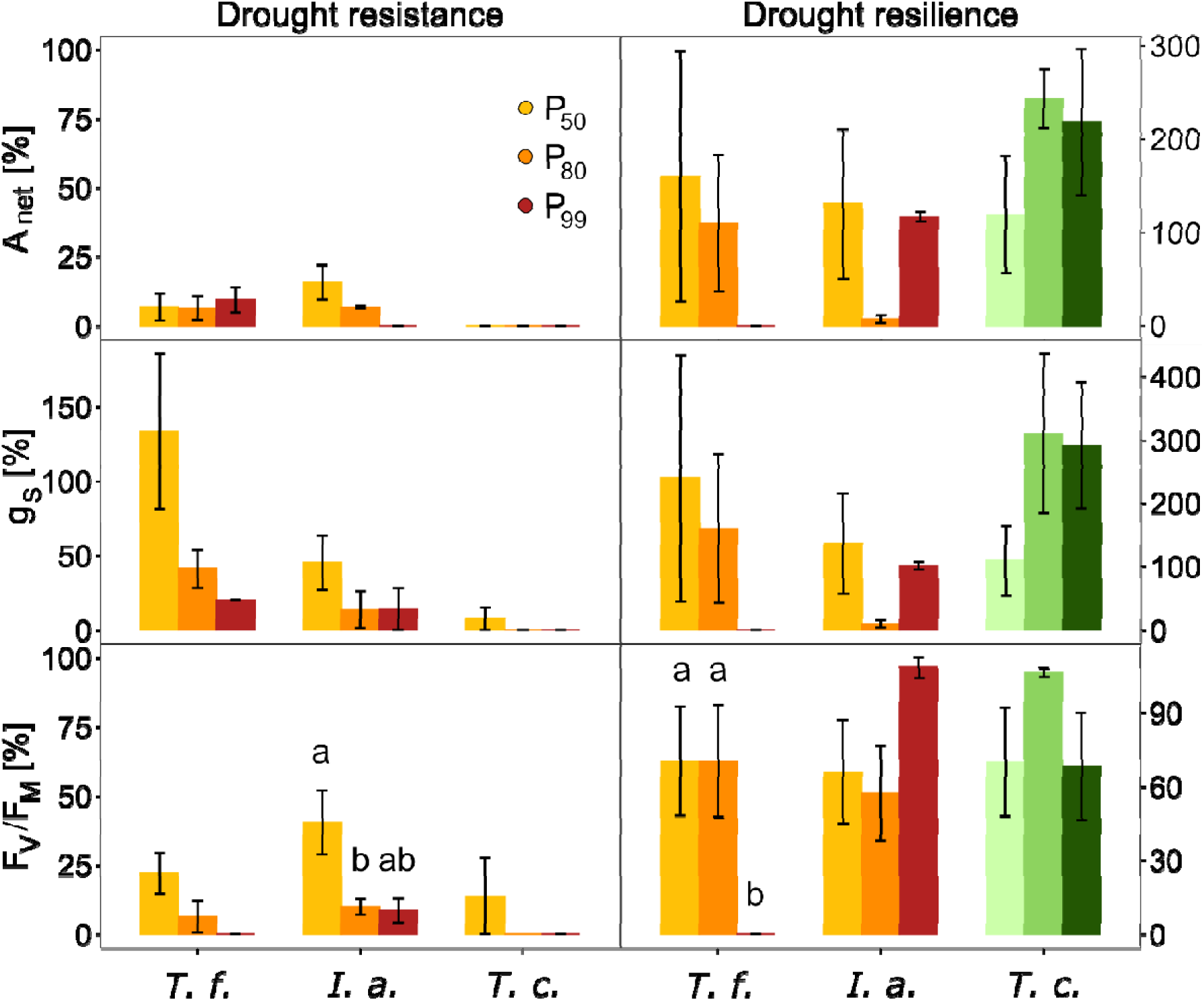
Drought resistance and resilience of the net CO_2_ assimilation (A_net_), stomatal conductance (g_S_), and chlorophyll fluorescence (F_V_/F_M_) for each species and drought treatment (*i*.*e*., P_50_, P_80_, and P_99_, corresponding to a loss of conductivity of 50, 80, and 99% at the leaf level, respectively) (means ± s.e., n = 3 individuals per species). Leaves that resprouted after the drought treatment are green. Different letters indicate significant differences between drought intensities (p < 0.05).

Interestingly, the individuals of *T*. *fortunei* exposed to the most severe drought (P_99_) lost all their leaves by the end of the recovery period, despite retaining some immediately after the drought (*i*.*e*., at T_1_) (Fig. S6). In contrast, *T*. *cordata* lost almost all of its leaves during the drought, but resprouted afterwards and exhibited higher A_net_ and g_s_ than the remaining leaves from other species (194 and 237% *vs*. 90.1 and 134% for *T*. *fortunei*, and 84.8 and 82.3% for *I*. *aquifolium*), although the difference was not statistically significant (Fig. 5 & Fig. S6). Resilience in F_v_/F_m_ was similar across all species but differed with drought severity. Plants subjected to the mildest drought (P_50_) displayed the highest resilience (25.4% compared to 5.46% for P_80_ and 2.86% for P_99_) (Fig. 5 & Fig. S6).

### 3.3 Predicted time to hydraulic failure

Simulations of THF indicated that all species reached stem P_99_ after a month under the tested experimental conditions (20.7°C mean T_air_), in agreement with field observations. THF values were 43.6, 33.9, and 34.1 days for *T*. *fortunei*, *I*. *aquifolium*, and *T*. *cordata, respectively* (Fig. 6). However, in *T*. *fortunei*, roots embolized faster than the leaves (n.s.), and significantly earlier than stems (*p* < 0.05) (25.1 and 43.6 *vs*. 86.9 days, respectively), revealing a strong organ-level variation (Fig. 6). A similar but less pronounced pattern was observed for *I*. *aquifolium*, where leaves and roots embolized faster than stems (33.9 and 43.0 *vs*. 66.1 days, *p* < 0.05). In contrast, all *T*. *cordata* organs were embolized in a span of 5 days (Fig. 6).

**Fig. 6:**
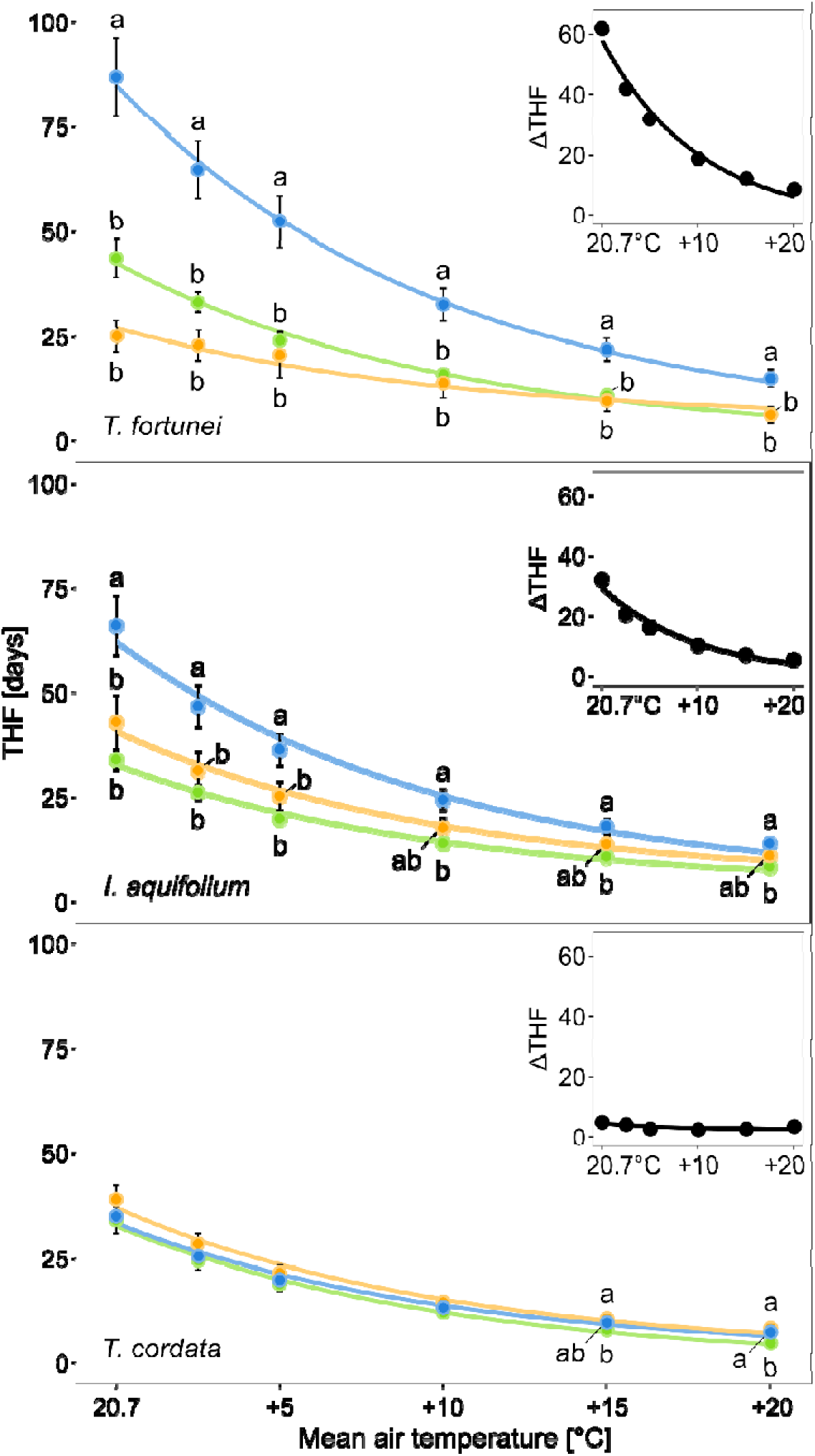
Simulated time to hydraulic failure (THF) in function of the mean air temperature of each organ (green = leaves; blue = stems; orange = roots) of the three species (means ± se, n = 30). We randomized the intraspecific physiological traits over 30 simulations, starting with the climatic data from the experiment (mean T_air_ = 20.7°C) up to +20 °C mean T_air_. Every relationship is significant (*p* < 0.05). Different letters indicate significant differences between organs at a given air temperature (p < 0.05). The top-right panels show the maximum difference in THF between organs in relation to the mean air temperature.

Increasing air temperature accelerated exponentially all organ hydraulic failures (R^2^ > 0.98) but did not alter the order in which organs reach hydraulic failure compared to our simulations at ambient temperature (Fig. 6). Still, the stem THF decreased more than the root and leaf THF in both *T*. *fortunei* and *I*. *aquifolium* (Fig. 6). This difference was driven by the depletion rate of the relative water content of the stems once disconnected from the roots and the leaves, that was much faster at higher temperatures than at ambient (Fig. S8). This was not happening in *T*. *cordata*, where all organs lost water simultaneously (Fig. S8).

## 4. Discussion

### 4.1 Embolism vulnerabilities of organs reflect anatomical and functional specialization in invasive palms

Palms were among the first plant groups in which xylem embolism was investigated (*e*.*g*., Sperry, 1985), yet surprisingly few recent studies have assessed their embolism resistance (Emilio *et al*., 2019). To our knowledge, this is the first study comparing hydraulic vulnerability across multiple palm organs, including roots.

We found that *T. fortunei* exhibited exceptionally drought-tolerant leaves and petioles, with leaf P_50_ values as negative as -7.44 MPa (Fig. 2), comparable to those reported in drought-adapted Australian conifers (Brodribb *et al*., 2017), and much more resistant than the typical values for tropical species (P_50_ ≈ -3 MPa) (Trueba *et al*., 2017; Smith-Martin *et al*., 2022). We observed a strong correlation between P_50_ and REL_50_ across all organs (Fig. 3), indicating that hydraulic failure is closely tied to the onset of cellular damage. This pattern likely reflects the progressive effects of drought-induced dehydration on membrane integrity (Mantova *et al*., 2023). The consistent ∼2 MPa gap between P_50_ and REL_50_ may be explained by vascular redundancies, where moderate xylem loss does not immediately compromise surrounding tissues (Sack *et al*., 2008; Mantova *et al*., 2023).

Our measurements of petiole xylem vulnerability in *T. fortunei* differ from those reported by Emilio *et al*. (2019), who couldn’t quantify P_99_ below -8 MPa and observed the first embolism events around -5.8 MPa. In contrast, we occasionally detected embolism events as early as - 2 MPa (Fig. S3), with considerable variation among individuals, potentially linked to differences in petiole diameter. This inter-individual variability suggests that embolism resistance in petioles may be influenced by structural traits and water storage capacity. Emilio *et al*. (2019) proposed that palms buffer xylem tension by mobilizing water from parenchyma tissues into the xylem, a mechanism also observed in tropical trees (Meinzer et al., 2008). The high water storage capacity of palm stems may help to prevent xylem embolism, which could be crucial for survival given the lack of secondary growth and limited repair mechanisms (Renninger and N. Phillips, 2016). Physiological measurements during the drought further support the role of water storage in the palm. Despite complete soil desiccation (Fig. 4), *T*. *fortunei* maintained high assimilation and stomatal conductance rates under dry conditions, particularly at P_50_, likely due to water storage depletion in the petiole parenchyma (Renninger and N. G. Phillips, 2016; Emilio *et al*., 2019). At the same time, F_v_/F_m_ decreased, indicating progressive tissue dehydration (Johnson, Jordan and Brodribb, 2018) (Fig. 4). These findings suggest that water storage in the palm buffered the effects of drought in the absence of resistance mechanisms such as stomatal closure or leaf shedding, which are present in the native species.

In stark contrast, *T. fortunei* roots exhibited low embolism resistance, with values comparable to those observed in tomato using microCT imaging (P_40_ = -1.38 MPa) (Skelton, Brodribb and Choat, 2017), but substantially less negative than those of other species such as wheat (P_50_ = -4.05 MPa) (Harrison Day and Brodribb, 2023) or olive trees (P_50_ = -7.10 MPa) (Rodriguez-Dominguez *et al*., 2018). The pronounced segmentation in hydraulic vulnerability between roots and leaves supports the concept of a drought protection hierarchy and likely reflects underlying anatomical differences, as previously observed between palm leaves and stems (Zimmermann and Sperry, 1983). Many monocots, including palms, feature tracheid-based xylem in their leaves and stems that tends to be effective at preventing embolism, while their roots rely predominantly on vessels (Carlquist, 2012; Emilio *et al*., 2019). This anatomical contrast likely underpines the pronounced variation in P_50_ we observed across organs in *T*. *fortunei* (Fig. 2), although more work would be required to confirm this. Root failure probably curtailed water uptake in P_99_ palms, which in turn prevented physiological restoration (Fig.L6) and ultimately triggered leaf loss within 45 days after the drought (Figs.L5 & S7).

The limited recovery potential observed in the invasive palm is consistent with findings by Funk and Zachary (2010), who used plant water status as a drought indicator and likewise found no greater recovery in invasive species compared to native ones. By contrast, studies reporting faster recovery of invasive species to drought often apply a fixed drought duration without accounting for physiological drought severity (*e*.*g*., Kelso, Wigginton and Grosholz, 2020; Zhang, Oduor and Liu, 2023), making comparisons difficult. In addition to their high drought vulnerability, roots also showed immediate cellular damage following embolism events (Fig. 2), suggesting low water storage to buffer embolism. In gymnosperms, high root water capacitance has been proposed as a compensatory mechanism for their generally low water transport efficiency, allowing daytime water demand to be met while root water stores are gradually replenished at night (McCarthy, Bourbia and Brodribb, 2025). By contrast, palms exhibit high transport efficiency due to their large xylem conduits (Olson *et al*., 2014), which may reduce their dependence on root water storage as carbon is allocated to transport tissues rather than storage. This trade-off between transport efficiency and storage capacity raises the question of how palms balance water supply and hydraulic risk across organs.

In sum, our findings highlight that *T. fortunei* exhibits strong hydraulic segmentation, with drought-tolerant leaves and stems supported by vulnerable roots. This organ-level pattern may reflect anatomical specialization and underscores the importance of integrating functional traits across organs when evaluating the drought resistance of invasive species.

### 4.2 Consequences of the palm hydraulic strategy on its invasion success

Contrary to expectations for invasive species, *T*. *fortunei* did not exhibit a typical resource- acquisitive strategy, which is often characterized by high A_net_ and g_s_ (Cavaleri and Sack, 2010; Liu *et al*., 2017; Montesinos, 2022). Instead, *T*. *fortunei*’s functional traits resembled those of native evergreen *Ilex aquifolium* and both retained their leaves for extended periods under drought conditions (Fig. 4). Leaf persistence in evergreen species has been associated with vulnerability segmentation between organs (*e*.*g*., Jin, Wang and Zhou, 2019), as observed in our data. Contrastingly, the deciduous *T*. *cordata* shed most leaves at relatively moderate water potentials (around -2 MPa), just before the onset of major xylem dysfunction (P₅₀ = -2.41 MPa). This early leaf shedding likely serves as an adaptive mechanism to reduce transpirational water loss and protects more vital organs from embolism (Wolfe, Sperry and Kursar, 2016). While defoliation incurs carbon costs, resprouted leaves in *T*. *cordata* showed photosynthetic rates up to four times higher than pre-drought levels (Fig. 4), possibly due to resource investments in the rhizosphere during the drought (Hagedorn *et al*., 2016) and increased Huber value following canopy reduction, similarly as observed in stand thinning experiments (Han, 2012). Such dynamic plasticity in leaf turnover and resource allocation may represent an important drought resilience strategy in deciduous species, contrasting with the more conservative and embolism-resistant behavior observed in evergreen ones.

Previous studies in natural forests have similarly reported lower or comparable assimilation rates for *T*. *fortunei* relative to co-occurring native species (Juillard *et al*., 2024). Given that its respiratory rates are comparable or smaller than native species (Juillard *et al*., 2025), these findings suggest that its invasive success may instead rely on traits associated with low carbon costs and high resource investment in reproductive output (*e*.*g*., propagule pressure), rather than on superior resource acquisition. This contrasts with many annual invasive plants, which often display high assimilation rates combined with low water-use efficiency, enabling rapid post-drought recovery consistent with a resource-acquisitive strategy. However, drought recovery in long-lived invasive species remains understudied. Emerging evidence suggests that such species may face a trade-off between water-use efficiency and growth (Sanders *et al*., 2024), indicating that success in dry environments may favor conservative hydraulic traits. Nonetheless, this hypothesis remains largely untested.

Palms originated during the warm and humid Cretaceous (Reichgelt, West and Greenwood, 2018), and may have evolved for optimised water transport efficiency to aboveground organs at the expense of root embolism resistance. Embolism resistance comes at a carbon cost and limits water transport capacity (Pittermann, 2010), conferring an important plant hydraulic trade-off. A lack of strong selection pressure for root resistance to embolism in wet ancestral habitats may thus explain the priority for a high water transport efficiency. In accordance, our findings suggest that the spread of *T*. *fortunei* is limited to relatively moist environments, as intense drought events can induce irreversible root damage (Fig. 6). In Central Europe, naturalized palm populations are frequently found in humid microhabitats such as lakeshores and riparian zones (Berger and Walther, 2006), though a direct causal link between water availability and its distribution has yet to be firmly established (Conedera *et al*., 2018).

Our data indicate that even mild drought could compromise root integrity despite apparent xylem resistance in the leaves, potentially reducing the palm’s capacity for recovery. In the future, elevated temperatures could accelerate dehydration across all organs (Fig. 6), compounding drought stress. It is plausible that mortality thresholds in palms are determined by embolism in the meristematic tissues, which are critical zones for leaf and root regeneration and may embolize later than other organs. Understanding this threshold will be crucial for predicting the long-term drought resilience and invasive potential of palm species under climate change.

## Acknowledgments

This study was supported by the Sandoz Family Foundation. CG and CB were supported by the Swiss National Science Foundation (310030_204697, CRSK-3_220989). We thank Alex Tunas Corzon, Patrick Favre, Quentin Gaillard, Laura Mekarni, Martin Metier, Arianna Milano, Alvaro Poretti, and Jolan Wicht for their support during the experiment setup and data analysis.

## Author contributions

All authors designed the study. TJ, KJ, TB, and MB contributed to the methodology. TJ and MB performed the experiments. TJ analyzed data a wrote the first version of the manuscript, with input from all authors. All authors approved the final manuscript.

## Data availability

All data supporting the findings of this study will be made openly available in Envidat.

