## Supplementary Material for "Root vulnerability to embolism and lack of physiological recovery limit the competitive ability of an invasive palm"

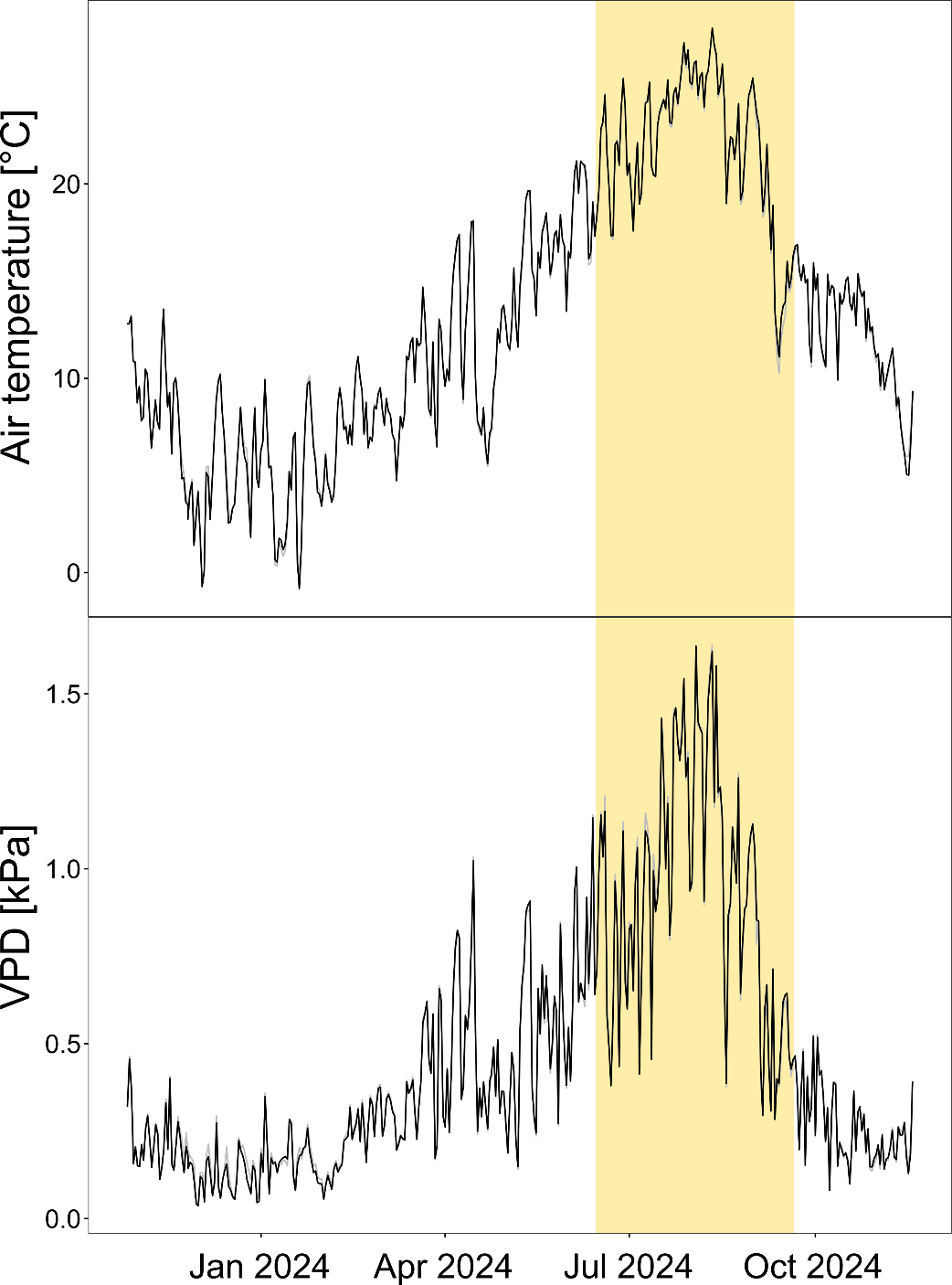

### Supplementary Material

#### **Fig. S1**: Daily air temperature (top) and vapor pressure deficit (VPD; bottom) inside the two greenhouses from November 2023 to November 2024. The drought and rewatering period (June-September 2024) is highlighted in yellow.

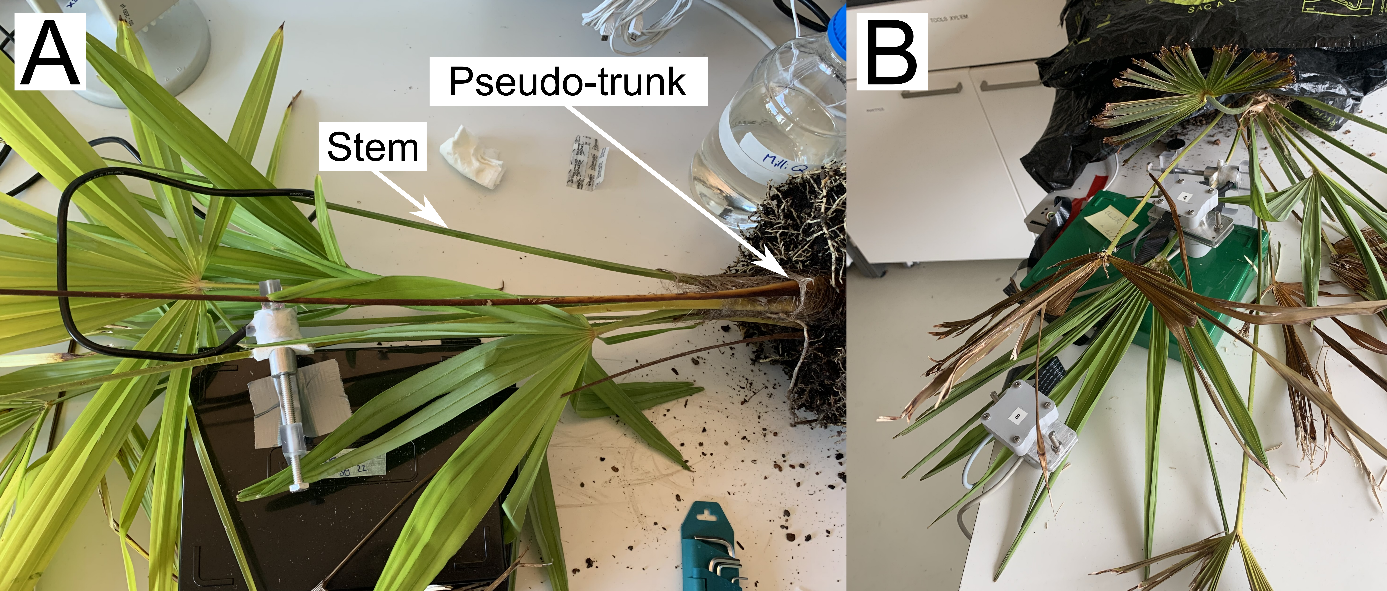

#### **Fig. S2**: (A) Photograph of a psychrometer on the petiole of *T*. *fortunei* (*i*.*e*., what we considered as a stem for the palm, and not the pseudo-trunk) and (B) cavicams on the leaves and stems of the palm.

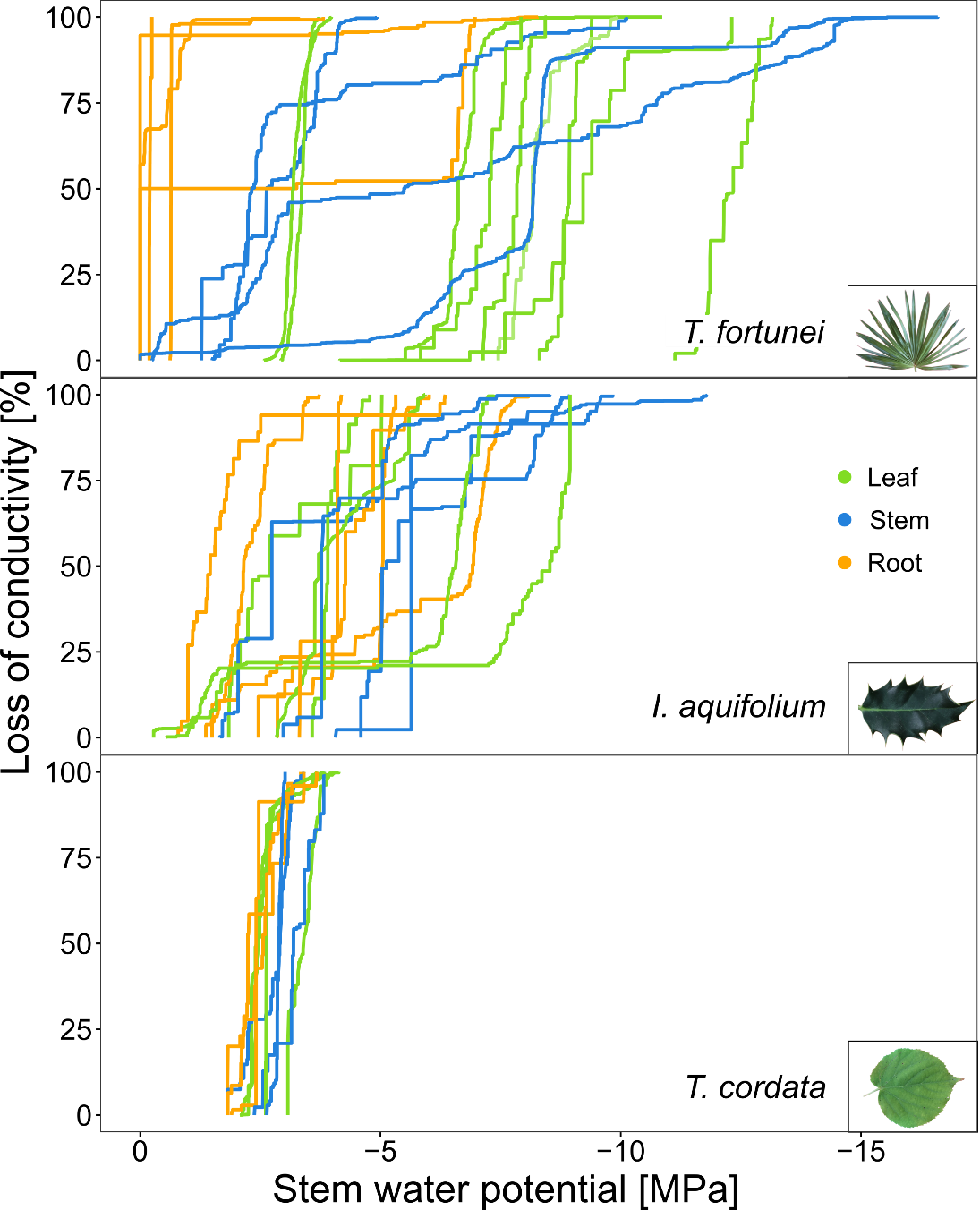

#### Fig. S3: Stem water potential in relation to embolism measured with the optical method for each organ (green = leaves; blue = stems; brown = roots) in each individual (n = 3-9). For *T*. *fortunei*, the petiole was measured instead of the stem due to technical limitations.

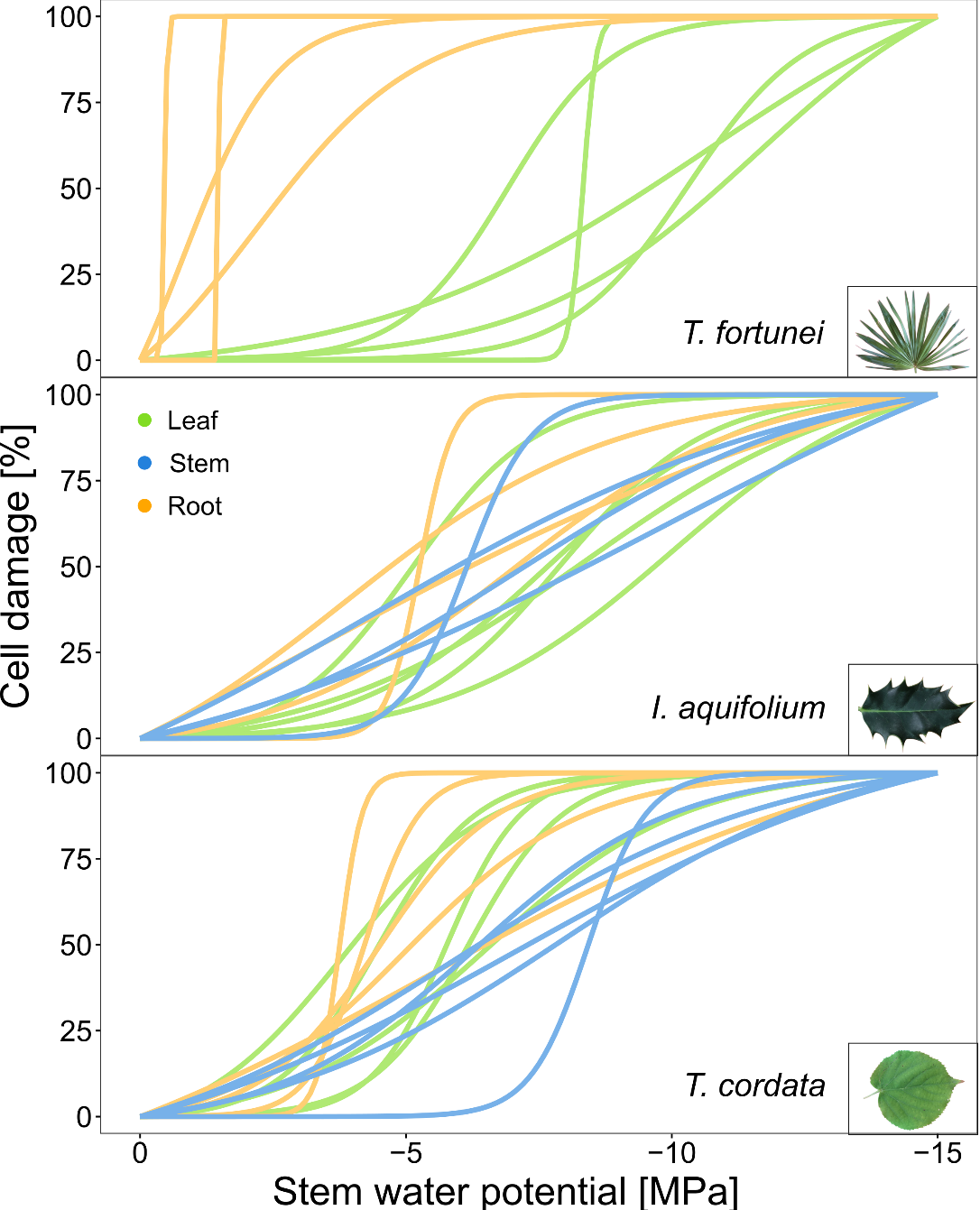

#### Fig. S4: Stem water potential in relation to cell damage for each organ (green = leaves; blue = stems; brown = roots) of each plant. Each curve was fitted with a logistic curve from 12-18 measurements. We could not measure petioles in *T*. *fortunei* as it requires cutting the complete leaflet

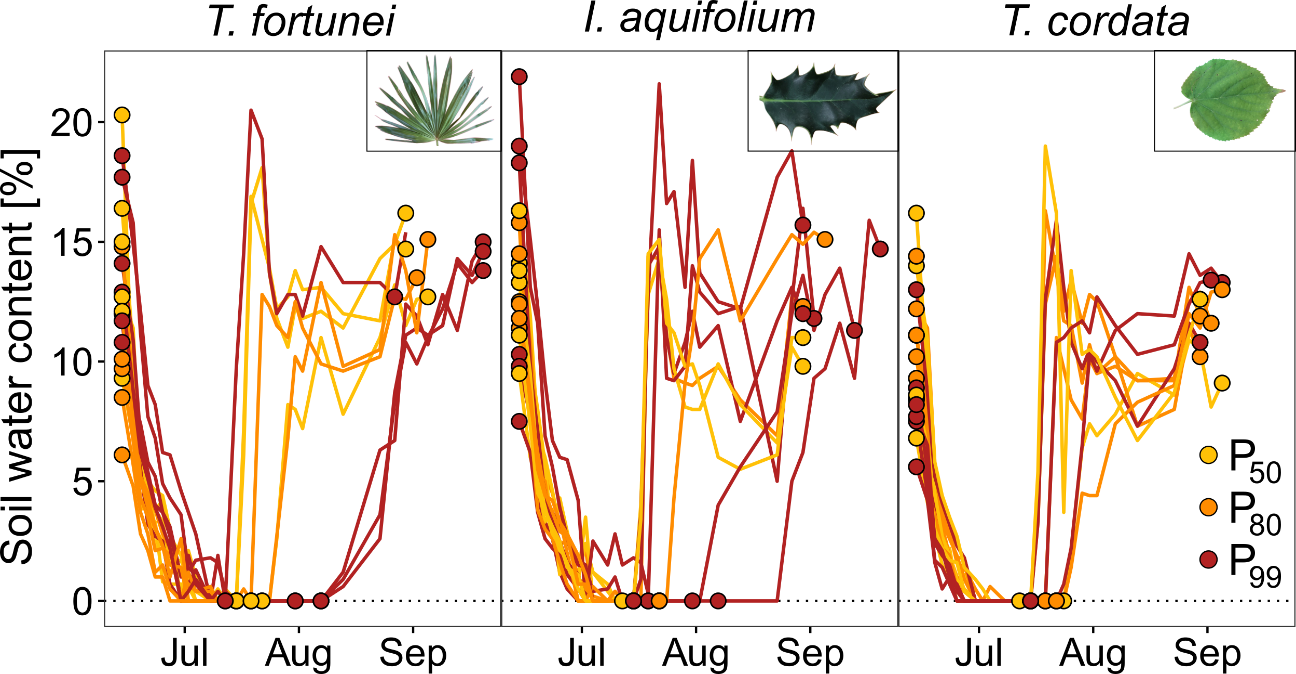

#### Fig. S5: Soil water content over time for each individual of the three species. Colors represent the different drought treatments, inducing a loss of conductivity of 50, 80, and 99% (P50, P80, and P99, respectively) at the leaf level.

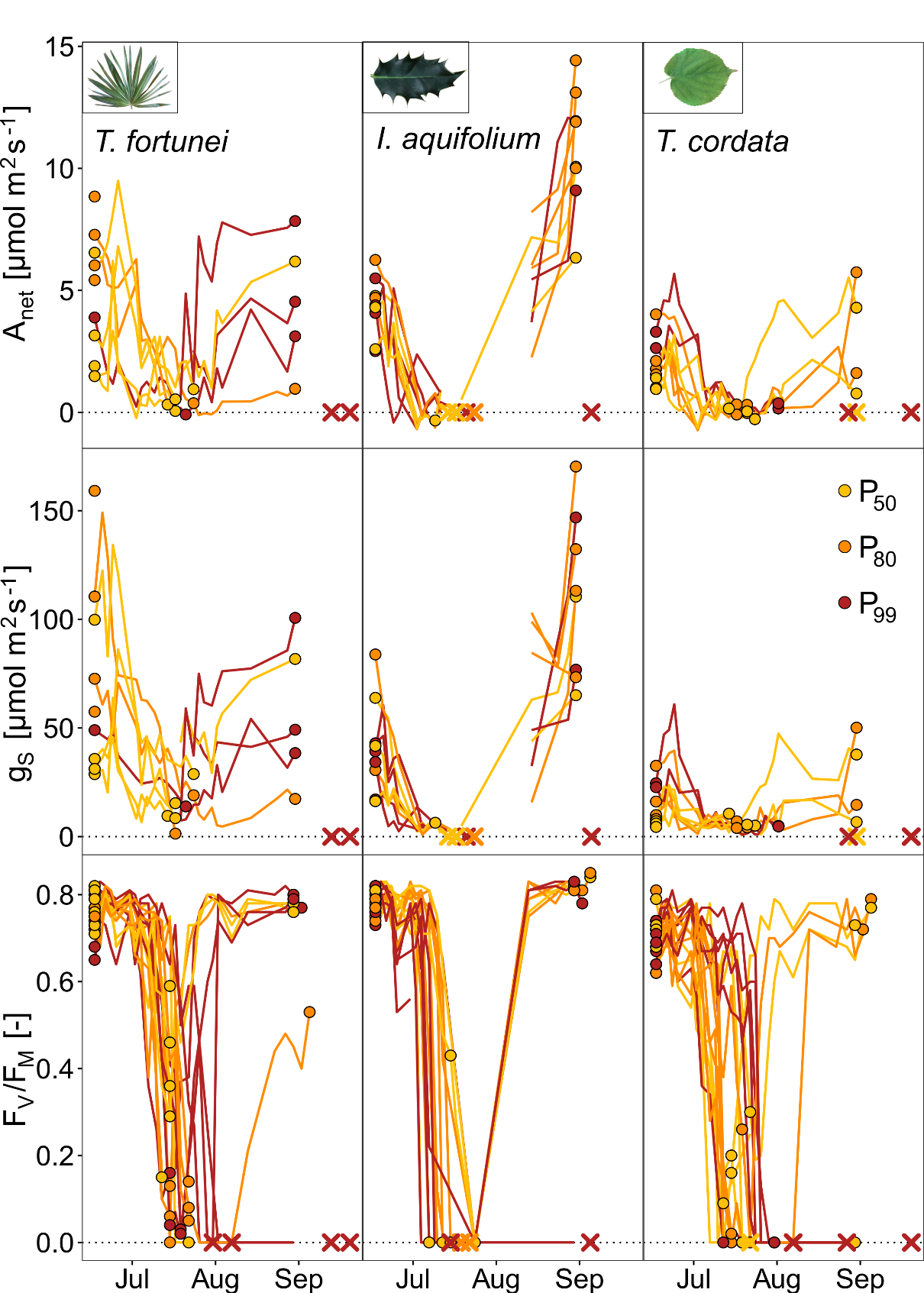

#### **Fig. S6:** Net CO_2_ assimilation (A_net_), stomatal conductance (g_s_), and chlorophyll fluorescence (F_v_/F_m_) over time for each individual of the three species. Colors represent the different drought treatments inducing a loss of conductivity of 50, 80, and 99% (P_50_, P_80_, and P_99_, respectively) at the leaf level.

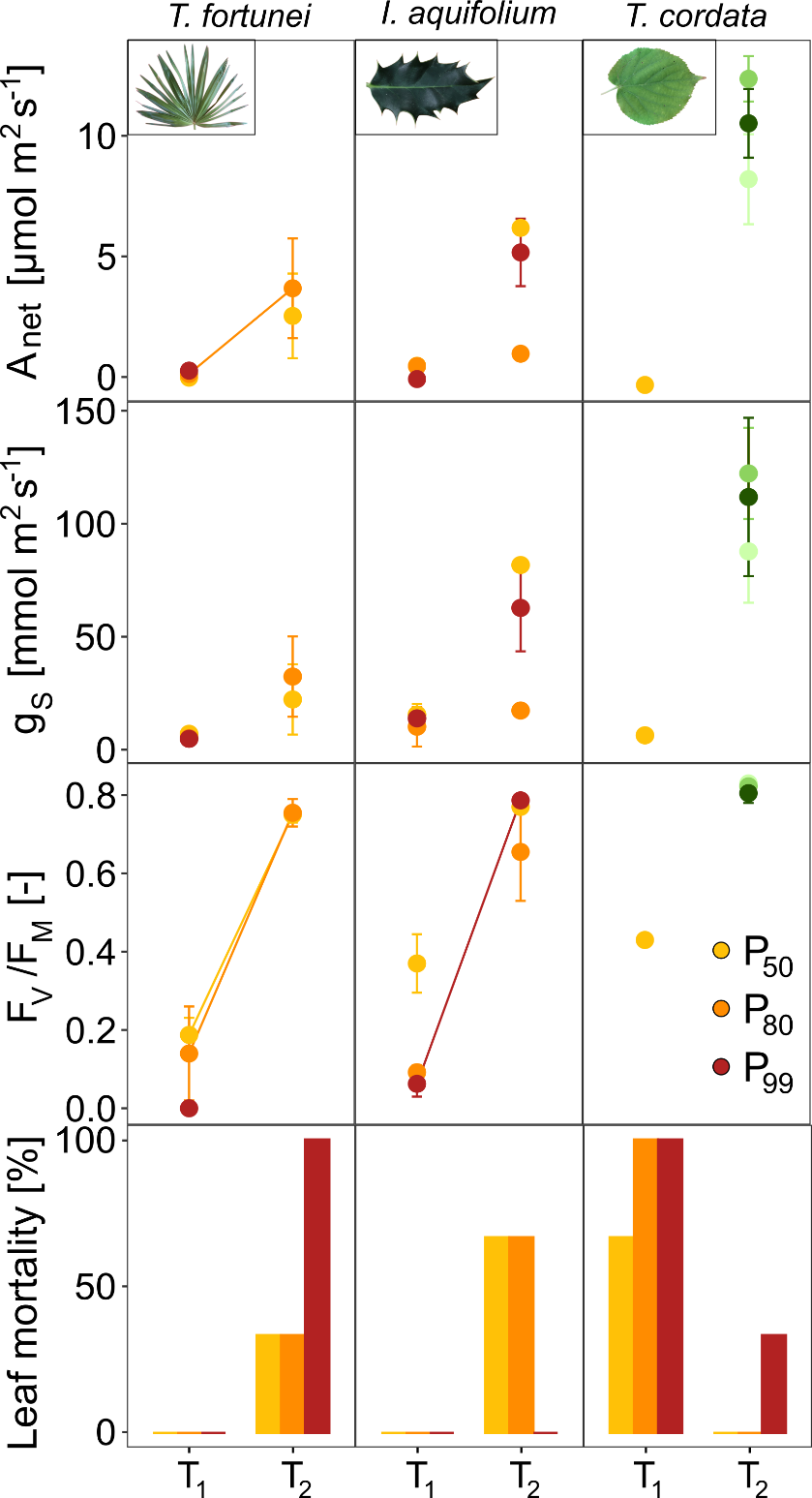

#### Fig. S7: Net assimilation (A_net_), stomatal conductance (g_s_), chlorophyll fluorescence (F_V_/F_M_), and leaf mortality (means +/- s.e., n = 3 individuals per species) at the peak of the drought treatment (T_1_), and after 45 days of rewatering (T_2_) for each species. Colors represent the different drought treatments inducing a leaf embolism of 50, 80, and 99% (P_50,_ P_80_, and P_99_, respectively). Significant differences between T_1_ and T_2_ are highlighted with a line (*p* < 0.05). Leaves that resprouted after the drought treatment are shown in green.

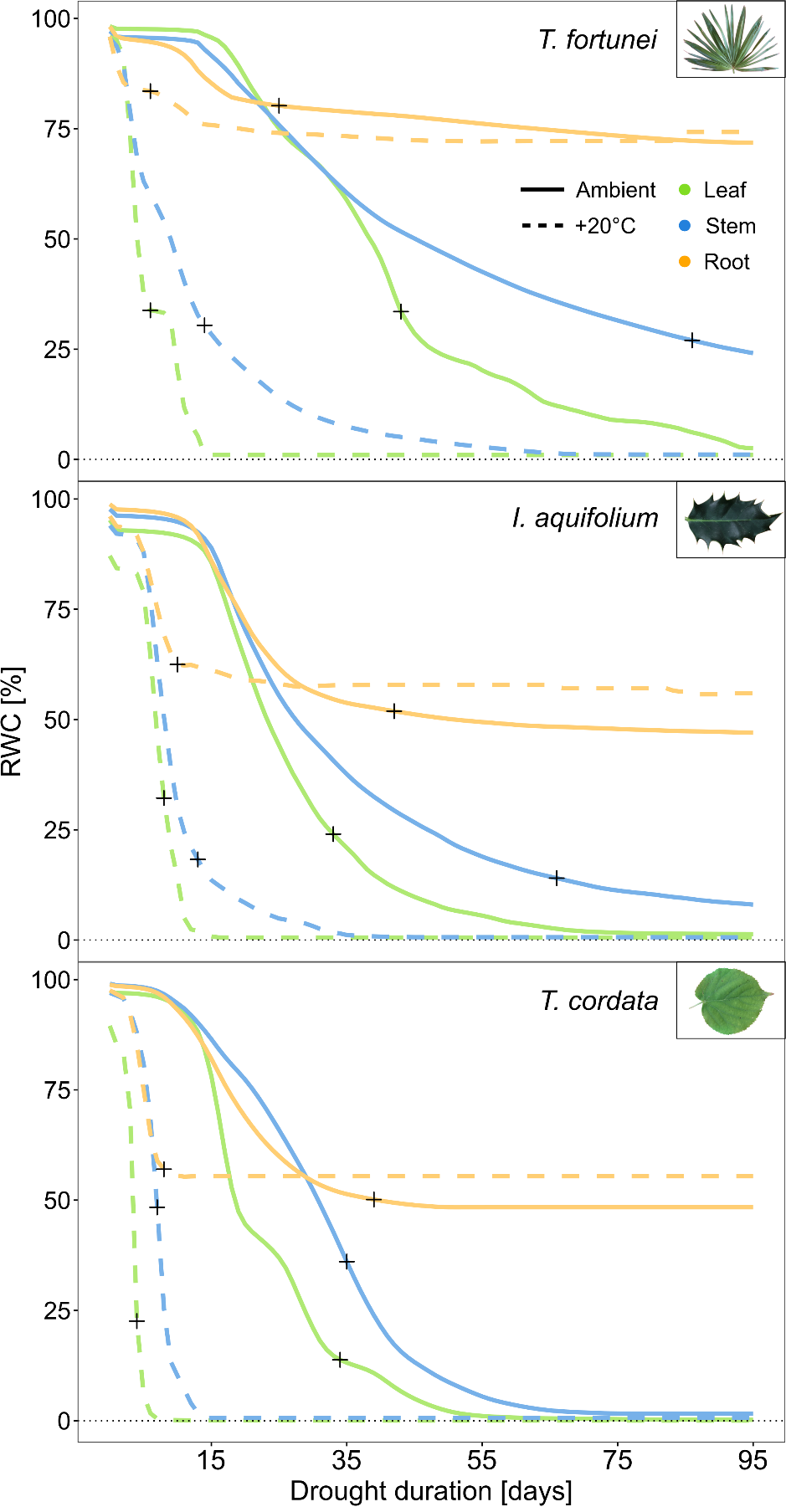

#### Fig. S8: Simulated relative water content (RWC) in relation to drought duration at ambient (20.7°C, solid line) and high mean air temperature (+20°C, dashed line) for each species and organs (n = 30 individuals). Black crosses indicate the time to mean hydraulic failure of each organ and species. Simulations were done with the SurEau model.

| **Variable** | **Unit** | **Value** | | | **Measurements** |
| --- | --- | --- | --- | --- | --- |
| **Environment** | | | | | |
| Latitude | ° | 46 | | |  |
| Day of year | - | 165 | | |  |
| Min. air temperature | °C | 15 | | |  |
| Max. air temperature | °C | 25 | | |  |
| RH min | % | 50 | | |  |
| RH max | % | 85 | | |  |
| PAR max | µmol | 1200 | | |  |
| Wind | m s^-1^ | 2 | | |  |
| Soil type | - | Clay Loam | | |  |
| Soil depth | m | 0.23 | | | Measured |
| Soil width | m | 0.2 | | | Measured |
| **Species traits** | | | | | |
|  |  | *T. fortunei* | *I. aquifolium* | *T.cordata* |  |
| Total height | m | 0.6 | 0.65 | 1.07 | Measured |
| Canopy height | m | 0.5 | 0.5 | 0.5 | Measured |
| Canopy width | m | 0.3 | 0.3 | 0.3 | Measured |
| Leaf size | mm | 10 | 40 | 60 | Measured |
| DBH | cm | 2.24 | 1.28 | 1.43 | Measured |
| LAI | m^2^ m^-2^ | 2.4 | 1.5 | 3.9 | Measured |
| Leaf succulence | g m^-2^ | 187.1 | 367.9 | 62.1 | Measured |
| **Stomata** | | | | | |
| g_min_ | mmol m^-2^ s^-1^ | 3.84 | 2.76 | 5.98 | Measured |
| g_max_ | mmol m^-2^ s^-1^ | 120 | 100 | 130 | Measured |
| P_g_12_ | MPa | -1 | -2 | -0.5 | Estimated |
| P_g_88_ | MPa | -2 | -4 | -1.5 | Measured |
| **Hydraulics** | | | | | |
| P50, leaf | MPa | -7.44 | -5.09 | -2.74 | Measured |
| Slope at P50, leaf | MPa | 166.5 | 100 | 150 | Measured |
| P50, stem | MPa | -4.66 | -5.73 | -3.00 | Measured |
| Slope at P50, stem | MPa | 14.78 | 10 | 250 | Measured |
| P50, roots | MPa | -1.33 | -4.00 | -2.41 | Measured |
| Slope at P50, roots | MPa | 1660 | 150 | 100 | Measured |
| Ψ_TLP_ | MPa | -3 | -2.5 | -1.9 | Measured |
| Elasticity modulus (ε) | MPa | 25 | 9.5 | 16 | Measured |

#### Table S1: Main input parameters used in the SurEau model
